# Superior *ab initio* Identification, Annotation and Characterisation of TEs and Segmental Duplications from Genome Assemblies

**DOI:** 10.1101/190694

**Authors:** Lu Zeng, R. Daniel Kortschak, Joy M. Raison, Terry Bertozzi, David L. Adelson

## Abstract

Transposable Elements (TEs) are mobile DNA sequences that make up significant fractions of amniote genomes. However, they are difficult to detect and annotate *ab initio* because of their variable features, lengths and clade-specific variants. We have addressed this problem by refining and developing a Comprehensive *ab initio* Repeat Pipeline (CARP) to identify and cluster TEs and other repetitive sequences in genome assemblies. The pipeline begins with a pairwise alignment using krishna, a custom aligner. Single linkage clustering is then carried out to produce families of repetitive elements. Consensus
sequences are then filtered for protein coding genes and then annotated using Repbase and a custom library of retrovirus and reverse transcriptase sequences. This process yields three types of family: fully annotated, partially annotated and unannotated. Fully annotated families reflect recently diverged/young known TEs present in Repbase. The remaining two types of families contain a mixture of novel TEs and segmental duplications. These can be resolved by aligning these consensus sequences back to the genome to assess copy number vs. length distribution. Our pipeline has three significant advantages compared to other methods for *ab initio* repeat identification: 1) we generate not only consensus sequences, but keep the genomic intervals for the original aligned sequences, allowing straightforward analysis of evolutionary dynamics, 2) consensus sequences represent low-divergence, recently/currently active TE families, 3) segmental duplications are annotated as a useful by-product. We have compared our *ab initio* repeat annotations for 7 genome assemblies (1 unpublished) to other methods and demonstrate that CARP compares favourably with RepeatModeler, the most widely used repeat annotation package.

**Author summary:** Transposable elements (TEs) are interspersed repetitive DNA sequences, also known as ‘jumping genes’, because of their ability to replicate in to new genomic locations. TEs account for a significant proportion of all eukaryotic genomes. Previous studies have found that TE insertions have contributed to new genes, coding sequences and regulatory regions. They also play an important role in genome evolution. Therefore, we developed a novel, *ab initio* approach for identifying and annotating repetitive elements. The idea is simple: define a “repeat” as any sequence that occurs at least twice in the genome. Our *ab initio* method is able to identify species-specific TEs with high sensitivity and accuracy including both TEs and segmental duplications. Because of the high degree of sequence identity used in our method, the TEs we find are less diverged and may still be active. We also retain all the information that links identified repeat consensus sequences to their genome intervals, permiting direct evolutionary analysis of the TE families we identify.

## Introduction

Thousands of genomes have been sequenced thanks to decreased cost and increased speed of DNA sequencing methods. The explosion of genome sequences has expanded our knowledge of repetitive DNA, which is an important component of the genomes of almost all eukaryotes. Repetitive DNA is made up of sequences that have been duplicated. Some repetitive elements are able to replicate to new genomic locations and are referred to as transposable elements (TEs). TEs are known to account for a significant proportion of genome sequences in eukaryotes, varying from a few percent to the majority of the genome. For example, around 50% of the human [1] and 85% of the maize genome are TEs [2]. Therefore, it is important to have an efficient and accurate *ab initio* method of identifying and annotating repeats in newly sequenced genomes.

Repetitive DNA sequences can be divided into three major categories: tandem repeats, segmental duplications and transposable elements. Tandem repeats are repeated DNA sequences that are directly adjacent to each other and account for 3% of the human genome [3].

Segmental duplications (SDs, also termed “low-copy repeats”) are DNA sequences of variable sequence length (ranging from 1kb to 400kb) and a high level of sequence identity. SDs are identified from pairwise local alignments generated with BLAST using arbitrary criteria (>90%id, >1000bp length) [4]. Because SD identification is based on local alignments, repeat masked genome sequences are used as input to remove the enormous number of alignments produced by TEs that would overwhelm the SD output. This means that repeat identification and annotation is currently required before SDs can be identified.

Transposable elements are the most prevalent repetitive sequences in eukaryotic genomes, and fall into two major classes: those moving via direct cut and paste of their DNA sequences (DNA transposons) and those moving/replicating via a copy and paste mechanism with an RNA intermediate (retrotransposons). DNA transposons encode a transposase gene that is flanked by two *Terminal Inverted Repeats* (TIRs) [5]. The transposase recognizes these TIRs to excise the transposon DNA, which is then inserted into a new genomic location by cut and paste mobilization [6].

Retrotransposons can be subdivided into two groups: those with long terminal repeats (LTRs), and those without LTRs (non-LTR). Endogenous retroviruses (ERVs) are domesticated remnants of retroviral infection and full-length ERVs encode an array of proteins (*gag*, *pol*, and *env*) flanked by LTRs [7]. The *env* protein allows ERVs to transfer to other organisms by infection [8] and thus ERVs can be acquired from the environment. LTR retrotransposons are the dominant retrotransposons in plants and are less abundant in mammals [9]. Similar to ERVs, LTR retrotransposons contain two long-terminal repeats that flank a 5-7kb long internal protein-coding domain [10] containing two open reading frames (ORFs): *gag* and *pol!*. The *gag* ORF encodes the structural protein that makes up a virus-like particle (VLP) [11]. The *pol* ORF encodes an enzyme needed for replication that contains protease (PR), integrase (IN), reverse transcriptase (RT), and RNase H (RH) domains required for reverse transcription and integration. Promoter and transcription termination signals are present in the LTRs that are divided into three functional areas: U3, R and U5. U3 contains the enhancer and promoter sequences that drive viral transcription [11]. However, due to the lack of *env* protein, LTR retrotransposons are not infectious; they are obligate intracellular elements [12].

Non-LTR retrotransposons include two sub-types: autonomous long interspersed elements (LINEs), and non-autonomous short interspersed elements (SINEs), that are dependent on LINEs for their replication [3]. Typical insertions of non-LTR retrotransposons are flanked by target site duplications, which result from micro-homology based repair during the insertion process [13].

LINEs contribute significantly to eukaryotic genomes. Full-length LINEs are around 6kb long and usually contain two ORFs flanked by 5′ and 3′ untranslated regions (UTRs). LINE 5′ UTRs possess an internal RNA polymerase II promoter, which allows them to be transcribed [1]. ORF1 can vary significantly from species to species, and can encode proteins with different characteristics [14]. ORF2 is similar across all LINEs and encodes a protein with endonuclease and reverse-transcriptase activities required for replication [14].

SINEs are much shorter; usually less than 500 base pairs. The 5′ region contains an internal RNA polymerase III promoter and the 3′ end contains an oligo dA-rich tail. *Alu* elements have no ORFs, therefore they have no coding capacity and are non-autonomous TEs. Because they share functional sequences at their 3′ with LINEs, they borrow the retrotransposition molecular machinery encoded by LINEs that bind to their 3′ end [1].

Repeats are computationally difficult to detect and annotate *ab initio* because of their abundance, varied features/sequence signatures, many length variants (truncated versions) and clade specificity. Many computational tools have been developed to detect TEs, and the most commonly used approaches can be divided into three categories:

1) Library-based methods (e.g. RepeatMasker [15]), that use sequence alignment to search a genome for homologs of known repeats from a database such as Repbase [16], Repbase is a manually curated repeat library of species-specific and pan-species TEs, and cannot be used to identify segmental duplications.
2) Signature-based methods, that rely on the fact that each class of TE has a set of unique sequence features such as target site duplications, a poly-A tail, terminal inverted repeats, etc… These methods search for the sequence signatures of the repeat class of interest (e.g. LTR_STRUC [17]). However, because repeat types are so varied, this method is usually only able to identify specific types of TE.
3) *Ab initio* consensus methods, four examples here are RepeatModeler (http://www.repeatmasker.org/RepeatModeler/), REPET [18], Red [19] and PILER [20]. RepeatModeler (RMD) is a *de novo* package that has been widely used for repeat identification and modeling that combines different programs: RepeatMasker, RepeatScout [21], RECON [22] and TRF (Tandem Repeat Finder) [23]. RepeatMasker identifies and masks interspersed repeats using curated libraries of consensus sequences supported by Dfam; Dfam contains entries corresponding to all Repbase TE entries, and each Dfam entry is represented by a profile hidden Markov model. RECON evaluates pair-wise similarities to build repeat consensus sequences. RepeatScout identifies and uses highly over-represented k-mers as seeds that are extended to produce multiple sequence alignments. However, RMD doesn′t identify the individual sequences used to derive the consensus sequences; making it impossible to confirm or assess the accuracy of the consensus sequences, or to directly analyse the repeat instances in the genome they are derived from.

Red is an *ab initio* tool for discovering repetitive elements in a genome. Red utilizes a Hidden Markov Model dependent on labeled training data, i.e. it is an instance of supervised learning. Red identifies candidate repetitive regions using adjusted counts of k-mers, score smoothing with a Gaussian mask and the second derivative test to find local maxima [19]. Red can detect both transposons and simple repeats. However, it only generates genome coordinates for repeats, without any annotation. Red output is therefore not useful for analysing repeat content or transposon evolution.

PILER can identify and cluster repeats based on pairwise whole-genome alignments. In contrast to previous methods that attempt to explain all the off-diagonal local alignments or hits, it focuses on identifying subsets of hits that form a pattern characteristic of a given type of repeat. PILER was orignally designed to use PALS to generate pairwise alignment; however, PALS cannot handle concurrent jobs and it was built for a 32-bit processor architecture, which makes it relatively time consuming and seriously limits PILER applicability to small genomes. Although any local aligner can be used to replace PALS, this requires attention to required alignment parameters, and hits need to be converted to PILER-compatible GFF format.

REPET is a package that requires a local aligner, three clustering tools (RECON, PILER and GROUPER [24]) and a knowledge/library based annotation pipeline [25]. REPET produces a very comprehensive output of repeat annotations, but excludes segmental duplications, is complex, requires genome annotation of gene models and is computationally expensive.

In order to address these limitations, we have created a comprehensive *ab initio* repeat pipeline (CARP) for identifying species-specific TE elements with high sensitivity and accuracy that deals with both TEs and segmental duplications. Our method also provides a full audit trail that links identified repeat sequences (and their genome intervals) to their families and consensus sequences. This permits direct evolutionary analysis of highly similar TE families.

## Methods

For a diagrammatic overview of our method for *de novo* discovery and annotation of repetitive elements from genome sequences see (Figure 1).

**Fig 1.**
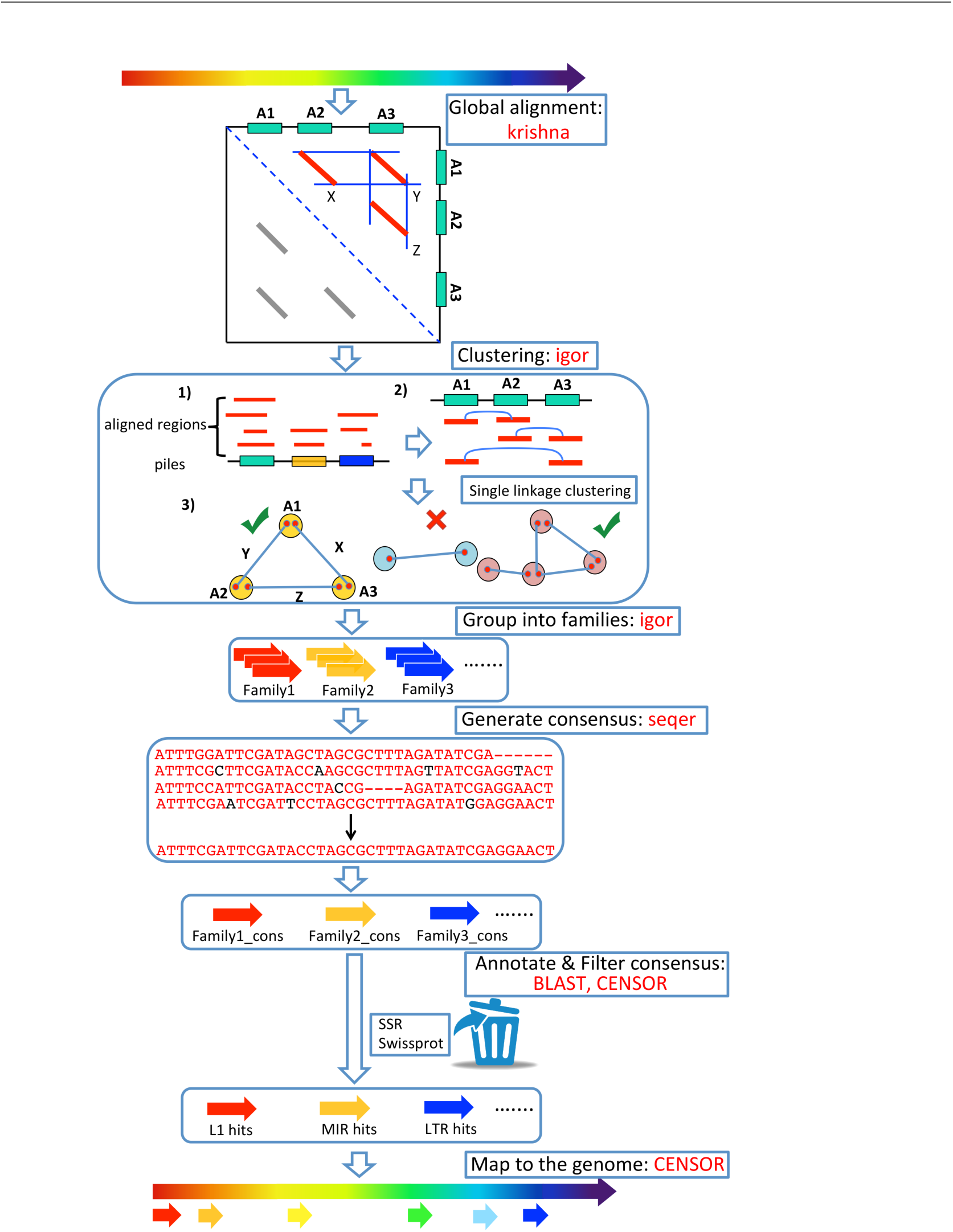
Comprehensive *Ab initio* Repeat Pipeline (CARP). Figure shows the detailed steps for CARP. Repetitive DNA is identified by all vs all pairwise alignment using krishna. Single linkage clustering is then carried out to produce families of repetitive sequences that are globally aligned to generate a consensus sequence for each family. Consensus sequences are filtered for non-TE protein coding genes and then annotated using Repbase and a custom library of retrovirus and reverse transcriptase sequences. The annotated consensus sequences are then used to annotate the genome. This is required to identify repeats with less than the threshold identity used for alignment that are overlooked during the initial pairwise alignment step.

### Datasets

Seven genomes were used in this study, 2 reptiles (anolis, *Anolis carolinensis* and bearded dragon, *Pogona vitticeps*), 1 bird (chicken, *Gallus gallus*), 2 monotremes (platypus, *Ornithorhynchus anatinus* and echidna, *Tachyglossus aculeatus*), 1 marsupial (opossum, *Monodelphis domestica*) and 1 eutherian mammalian (human, *Homo sapiens*). Most genomes are publicly available from the National Center for Biotechnology Information (NCBI). Supplementary table S1 lists the systematic name, common name, version, source and submitter for each genome assembly, and identifies privately acquired genomes. Supplementary table S2 shows the total genome sequence length and scaffold/contig N50 values, giving an approximation of the assembly quality. Supplementary table S3 compares the different sequencing technologies and methods.

### Comprehensive *ab initio* Repeat Pipeline (CARP)

Repeats were identified using a pipeline comprised of krishna/igor [26], MUSCLE (v3.8.31) [27] and WU-BLAST (v2.0) [28]. Krishna/igor is an improved version of PALS/PILER implemented in Go (https://golang.org/) that can find dispersed repeat families. A dispersed repeat family has members that are typically separated in the genome, i.e. that are rarely or never found in tandem, and are usually mobile elements such as retrotransposons [20]. Genome sequences were pairwise aligned using krishna (https://github.com/biogo/examples/krishna) with default parameters set at 94% sequence identity (-dpid) and a minimum alignment length (-dplen) of 250bp for most cases, except bearded dragon and chicken, which used ‐dpid 90% and ‐dplen 200bp. The resulting alignment intervals were then used as input for igor to define families of repeat sequences using the default parameters. Igor output was used as input for seqer to generate repeat consensus sequences for each cluster/family based on MUSCLE alignments. Only family members within 95% of the length of the longest family member were aligned, and to avoid consensus sequence expansion due to indels in the global alignment, a maximum of 100 randomly chosen sequences/family were included in the alignment. This process yielded three types of family: fully annotated, partially annotated and unannotated.

Identifiable repeat consensus sequences were annotated by using CENSOR [29] with the Repbase ‘Vertebrate’ library (downloaded on 1st March, 2016, includes 41,908 sequences). Further annotation of consensus sequences was based on WU-BLAST alignment against a comprehensive retroviral and retrotransposon protein database assembled from the NCBI [30], and against Swiss-Prot [31] to identify known protein-coding genes from large gene families inappropriately included in the repeat set. Consensus sequences identified as either simple sequence repeats (SSRs) or protein-coding sequences were removed from the consensus set. After acquiring all the annotated repeat consensus sequences, these annotated consensus sequences were then combined with the Repbase ‘Vertebrate’ library and CENSOR was used to annotate all repeat intervals in the source genome. Supplementary table S4 represents the summary of time consumed for each analysis step. For additional details of time consumption and memory use for each step, see Supplementary S1 Appendix.

## Method Evaluation

RepeatModeler (version 1.0.8) was used to evaluate the performance of CARP by applying it to the same seven genomes with default parameters, with WU-BLAST used as the alignment engine. A combination of the repeat consensus sequences generated by RepeatModeler and Repbase ‘Vertebrate’ library was also fed into CENSOR to annotate the repeat content for each genome.

### Identification Of Novel Repeat Sequences From Tested Genomes

In order to explore the unclassified consensus sequences generated by CARP, we extracted all unclassified repeat sequences from the seven genomes, and the R package ggplot2 was used to visualise their length distribution with respect to copy number.

For high copy number ( >2,000 copies), a coverage plot was used to investigate the positional distribution of genomic sequence fragments with respect to the unclassified consensus sequences. BLASTN and CENSOR were further used to characterise the consensus sequences from the coverage peaks of 5 unclassified consensus sequence examples found in the bearded dragon coverage plot.

Human (GRCh37) segmental duplication coordinates were also downloaded (http://humanparalogy.gs.washington.edu/build37/build37.htm) and BedTools [32] was used to merge the overlapping intervals from this data. We then used the human unclassified consensus sequences generated from both our *ab initio* method and RepeatModeler as libraries to run CENSOR against the merged segmental duplication data.

### Dendrogram construction from echidna nucleotide L2 sequences

Full-length echidna L2 consensus sequences (2~4kb) generated from CARP and RMD were extracted respectively, as well as the genome intervals that linked to the L2 consensus sequences from CARP. We then globally aligned the resulting sequences using MUSCLE (-maxiters 2). Alignments were trimmed with Gblocks [33] to remove large gaps (default parameters, allowed gap postions: with half). FastTree (v2.1.8) [34] was used to infer a maximum likelihood phylogeny from the global alignment, using a generalized time-reversible model (-gtr). Archaeopteryx v0.9901 beta was used to visualise the tree, including 166 genome intervals from CARP, 25 L2 consensus sequences from CARP and 4 consensus sequences from RMD.

### Classification of potentially active L2 elements

USEARCH [35] was then used to scan for open reading frames (ORFs) in those full-length L2 consensus sequences that were at least 60% of the expected length (⩾1.5kb nucleotide sequence for ORF2p, complete with start and stop codons and no inactivating mutations). After translation, ORF2p candidates were checked for similarity to known domains using HMM-HMM comparison [36] against the Pfam28.0 database [37] as of May 2015 (includes 16,230 families). ORF2p containing RT domains were extracted using the envelope coordinates from the HMMer domain hits table (-domtblout), with a minumum length of 200 amino acids.

## Results

### Consensus Generation

We identified and annotated repeats from seven genomes using both CARP and RepeatModeler. Because ancient transposable elements are highly diverged and already well described, we have implemented CARP to identify and annotate slightly diverged (recent) repetitive elements. CARP is based on whole-genome pairwise local alignment (default 94% identity), followed by clustering and consensus generation from clusters. This means all consensus sequences generated by CARP can be traced back to their input sequences and the original genomic sequence intervals of the input sequences. This provides an audit trail and the ability to easily carry out evolutionary and phylogenetic analysis of recently diverged, and hence recently active TEs. Because the initial clusters may contain gene families with many paralogs, we cleaned the consensus sequences by aligning them to Swiss-Prot and to a custom database assembled from retroviral and reverse transcriptase (RT) sequences from NCBI. We then removed consensus sequences that align to *bona fide* protein coding genes that do not annotate as retroviral/RT. Cleaned consensus sequences were then annotated with CENSOR using known TE reference sequences from Repbase. This resulted in three types of annotation: 1) well annotated, almost full length alignment to a Repbase reference sequence, 2) partially annotated, partial alignment with one or more Repbase sequences and 3) no significant alignment to a Repbase reference sequence. Partially annotated and unannotated consensus sequences were combined to produce the unclassified consensus repeat set.

CARP generated numerous consensus sequences (see Table 1), because TEs, particularly LINEs, are often 5^′^ truncated, generating many insertion length variants and because consensus generation is based on alignment pairs that are single-linkage clustered with a length constraint (within 95% of the longest family : member length). By comparing the repeat consensus sequences generated from CARP and RMD, we can see that CARP identified many more repeat sub-families, in contrast to RMD, which only generated a small number of broad consensus sequences. The latter are useful for masking, but are not as useful for studying TE evolution.

**Table 1.** Summary of consensus sequence libraries generated by CARP and RMD.

|  | Chicken |  | Bearded dragon |  | Anolis |  | Platypus |  | Echidna |  | Opossum |  | Human |  |
| --- | --- | --- | --- | --- | --- | --- | --- | --- | --- | --- | --- | --- | --- | --- |
|  | CARP | RMD | CARP | RMD | CARP | RMD | CARP | RMD | CARP | RMD | CARP | RMD | CARP | RMD |
| Well-annotated | 8,898 | 149 | 17,013 | 334 | 32,658 | 360 | 9,476 | 130 | 18,651 | 139 | 49,548 | 344 | 30,114 | 211 |
| Unclassified | 12,140 | 152 | 169,602 | 1080 | 45,201 | 1,361 | 165,231 | 466 | 72,245 | 454 | 25,221 | 768 | 23,199 | 542 |
| Total | 21,038 | 301 | 186,615 | 1414 | 77,859 | 1,721 | 174,707 | 596 | 90,896 | 593 | 74,769 | 1,112 | 53,313 | 753 |

### Consensus Classification

CARP generated annotated consensus sequences for all major TE types (except for SINEs in the chicken), whereas no SINE consensus sequences were produced by RepeatModeler in any of the species we tested (see Table 2). Based on this result, CARP was more sensitive for detecting SINEs compared to RMD. CARP generated many more consensus sequences than RMD and this is a function of the single linkage clustering used to identify families. Because many LINE insertions are 5’ truncated, leading to variable insertion sizes with a common 3′ end, the requirement for family members to be at least 95% as long as the longest family member means that many clusters are created across the insertion size continuum.

**Table 2.** Comparison of the total number of specific TE types in each method.

|  | Chicken |  | Bearded dragon |  | Anolis |  | Platypus |  | Echidna |  | Opossum |  | Human |  |
| --- | --- | --- | --- | --- | --- | --- | --- | --- | --- | --- | --- | --- | --- | --- |
|  | CARP | RMD | CARP | RMD | CARP | RMD | CARP | RMD | CARP | RMD | CARP | RMD | CARP | RMD |
| SINEs | 0 | 0 | 1,764 | 0 | 2,177 | 0 | 2,292 | 0 | 3,264 | 0 | 596 | 0 | 13,165 | 0 |
| LINE | 6,186 | 96 | 13,415 | 211 | 18,014 | 285 | 6,290 | 88 | 15,189 | 91 | 25,150 | 218 | 10,832 | 114 |
| LTR | 2,405 | 31 | 600 | 75 | 4,784 | 37 | 263 | 20 | 84 | 25 | 23,195 | 69 | 5,635 | 46 |
| DNA | 72 | 19 | 1,216 | 41 | 7,619 | 35 | 46 | 13 | 74 | 16 | 600 | 32 | 239 | 39 |
| Others | 235 | 3 | 18 | 7 | 64 | 3 | 585 | 9 | 40 | 7 | 7 | 25 | 243 | 12 |
| Total | 8,898 | 149 | 17,013 | 334 | 32,658 | 360 | 9,476 | 130 | 18,651 | 139 | 49,548 | 344 | 30,114 | 211 |

### Genome Repeat Content

CENSOR was used to annotate the repeat content in our data set of seven species because it uses minimal post-alignment processing of hits (see Table 3). In order to get a comprehensive annotation of repeats, we used a combination of the Repbase ‘Vertebrate’ library and repeat consensus sequences generated from CARP or RMD. Because CENSOR annotates based on the best hit, combining our consensus sequences with Repbase sequences allows annotation of genomic intervals most similar to either recent/less diverged repeats or Repbase repeats. As seen from Table 3, CARP performed consistently well in identifying and annotating repeats across all seven species (more detail in Supplmentary table S5~11).

**Table 3.** Comparison of repeat annotation for CARP and RMD. Summary of specific repeat content from CENSOR output, using a combined library of Repbase ‘Vertebrate’ with CARP or RMD consensus libraries. IR = Interspersed Repeats SD = Segmental Duplications

|  | Chicken |  | Bearded dragon |  | Anolis |  | Platypus |  | Echidna |  | Opossum |  | Human |  |
| --- | --- | --- | --- | --- | --- | --- | --- | --- | --- | --- | --- | --- | --- | --- |
|  | CARP | RMD | CARP | RMD | CARP | RMD | CARP | RMD | CARP | RMD | CARP | RMD | CARP | RMD |
| SINE | 0.09 | 0.12 | 1.72 | 1.68 | 4.02 | 4.14 | 19.51 | 19.32 | 19.86 | 19.56 | 10.44 | 10.46 | 11.25 | 11.34 |
| LINE | 7.73 | 7.67 | 11.61 | 11.72 | 14.65 | 14.78 | 20.40 | 20.39 | 25.68 | 25.76 | 28.21 | 28.68 | 18.88 | 18.93 |
| LTR | 3.37 | 3.46 | 2.38 | 3.03 | 5.98 | 5.86 | 1.34 | 1.50 | 1.15 | 1.34 | 10.62 | 10.64 | 9.13 | 9.46 |
| DNA | 2.60 | 2.27 | 3.55 | 3.54 | 12.84 | 12.90 | 1.72 | 1.69 | 1.67 | 1.45 | 3.06 | 2.95 | 4.49 | 4.50 |
| Other | 1.52 | 1.94 | 1.39 | 1.36 | 1.51 | 1.23 | 2.89 | 2.91 | 1.87 | 2.12 | 1.72 | 2.13 | 2.04 | 1.73 |
| IR | 15.31 | 15.46 | 20.65 | 21.33 | 39.00 | 38.91 | 45.86 | 45.81 | 50.23 | 50.23 | 54.05 | 54.86 | 45.79 | 45.96 |
| Potential SD | 1.99 | 0.69 | 22.22 | 13.82 | 12.02 | 8.28 | 12.60 | 1.07 | 5.96 | 0.92 | 3.93 | 1.76 | 2.94 | 0.19 |
| Total | 17.30 | 16.15 | 42.87 | 35.15 | 51.02 | 47.19 | 58.46 | 46.88 | 56.19 | 51.15 | 57.98 | 56.62 | 48.73 | 46.15 |

Compared to RMD, CARP identified approximately the same amount of sequence made up of interspersed repeats in all seven species. However, CARP identified far more of all seven genomes as derived from unclassified repeats. Because unclassified repeats are defined as not being classifiable using Repbase, these repeats must either be novel transposable elements, or repeated sequences that are not transposable. In Table 3 we have labeled the unclassified repeat contribution to the genomes as segmental duplications based on their properties (see below).

### Segmental Duplications

In order to determine if unclassified consensus sequences represent novel TEs or segmental duplications, we plotted the log10 transformed copy number of the unclassified CENSOR hits against their log10 transformed length for all seven genomes (see Figure 2). For both RMD and CARP unclassified sequences in all seven species, copy number increased with length, as determined by the regression line (Supplementary table S12). However, CARP unclassified sequences were generally present at much lower copy number, a strong indication of segmental duplication. The small number of high copy number (>2000) CARP unclassified sequences were examined for the presence of either novel TEs or partial TEs.

**Fig 2.**
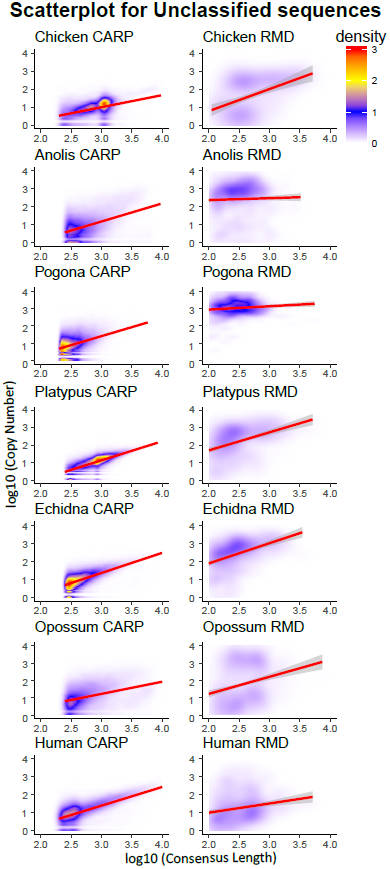
Scatter plot of unclassified sequence copy number *versus* length. Figure plots the copy number of unclassified sequences annotated using CENSOR and combined libraries, with respect to their length. Both copy number and length were log10 transformed. Red regions on the plot indicate high density, while blue regions indicate low density. Linear regression lines are plotted in red, with STANDARD ERROR represented by the gray shadow around the lines.

In order to determine if the small number of unclassified CENSOR hits with copy numbers >2000 were novel or partially annotated TEs, we used coverage plots for the CARP unclassified consensus sequences to look for high copy number subsequences with TE properties. Figure 3 shows the top 5 high copy number CARP unclassified consensus sequences from bearded dragon as an example. BLASTN and CENSOR annotations were also used to characterize these consensus sequences in terms of TE or gene model homology. From Figure 3 we can see that coverage plots for high copy number CARP unclassified CENSOR hits were of two types: those incorporating high copy subsequences (Figure 3A,C,E) and those with uniform high coverage (Figure 3B,D). Close examination of the high copy subsequences from Figure 3A,C,E show that known TE annotation cannot explain the high copy number subsequences detected in these families. Because these three consensus sequences were derived from families with a small number of members, the observed high copy subsequences may indicate similarity to unclassified TEs or TE fragments that are present as part of a small number of highly conserved segmental duplications. For the uniform high coverage family 0309690 (Figure 3B), CENSOR annotated one end as the 5′ end of a DNA transposon (Mariner-3N1), and the other end as the 3′ end of the same DNA transposon, likely indicating a novel variant of a known DNA transposon. For the uniform high coverage family 137078 (Figure 3D), there is no known TE annotation, only annotation for a part of GPR34, a probable G-protein coupled receptor gene.

**Fig 3.**
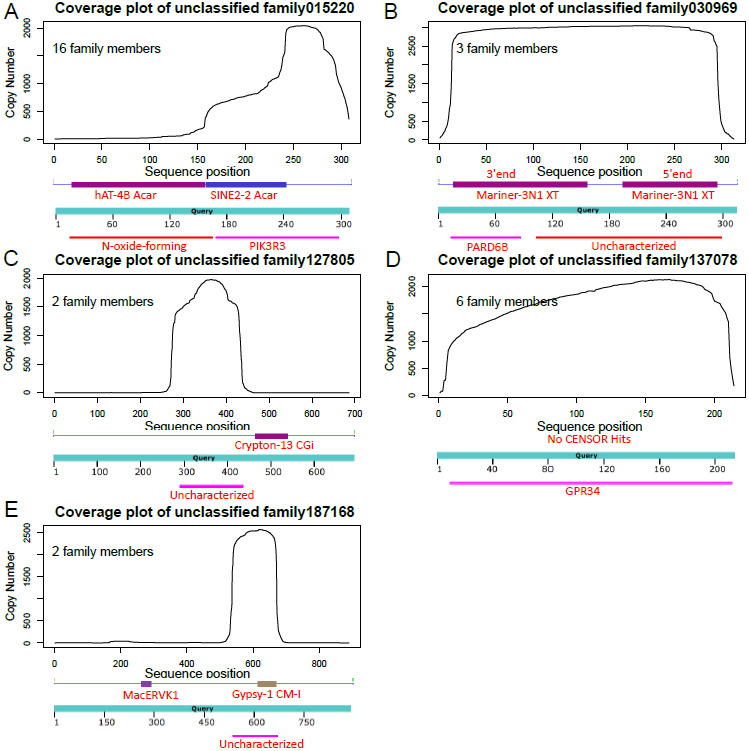
Coverage plot of the top 5 high copy CARP unclassified consensus sequences from the bearded dragon. A) CENSOR and BLASTN annotation of the peak coverage region in unclassified family 015220; B) CENSOR and BLASTN annotation of the peak coverage region in unclassified family 0309690; C) CENSOR and BLASTN annotation of the peak coverage region in unclassified family 127805; D) CENSOR and BLASTN annotation of the peak coverage region in unclassified family 137078; E) CENSOR and BLASTN annotation of the peak coverage region in unclassified family 187168. The number of family members identified by krishna/igor used for consensus sequence generation is shown in the upper left corner of each panel.

Based on the above results, we conclude that the vast majority of unclassified consensus sequences represent segmental duplications. We have therefore labeled these annotations accordingly in Table 3. In our final annotation, significant fractions of the genomes from our seven test species were annotated as SD, particularly in bearded dragon (24.68%), anolis (12.02%) and echidna (12.60%) (see Table 3).

Because the human genome has the best SD annotation of our seven species, we compared segmental duplication coordinates downloaded from the human ‘Segmental Duplication Database’ to our CARP unclassified CENSOR hits. Approximately 70% of human SD overlapped with CENSOR hits from CARP unclassified consensus sequences, confirming our conclusion above. Only 30% of human SD overlapped with RMD unclassified CENSOR hits.

### CARP classification of TEs allows insight into TE evolutionary dynamics

Because CARP enabled us to identify and classify recently diverged repeats, we were able to determine whether those repeats were consistent with recent TE activity/family expansion. We used the echidna to illustrate this, as this is the first repeat identification and annotation of the echidna genome. L2 and its non-autonomous SINE companion, mammalian-wide interspersed repeat (MIR, MON-1 in monotremes), are the most abundant and active repeats in monotremes (see Supplementary Table S9). This is in contrast to metatheria and eutheria (marsupials and placentals) where they are inactive due to extinction 60-100 Myr ago.

L2s were defined as potentially active if they contained an intact ORF2 (regardless of the state of ORF1), as this meant that they were capable of either autonomous retrotransposition [38] and/or mobilisation of SINEs [39]. CARP identified numerous long L2 elements (2~4kb) in the echidna genome.

More than 66% (110/166) of these were potentially active based on the above criteria (Figure 4A) and some clusters of potentially active elements at the tips of short branches, were consistent with “hot” or hyperactive elements. This differed significantly from the RMD result, which generated only four long consensus sequences (Figure 4C).

**Fig 4.**
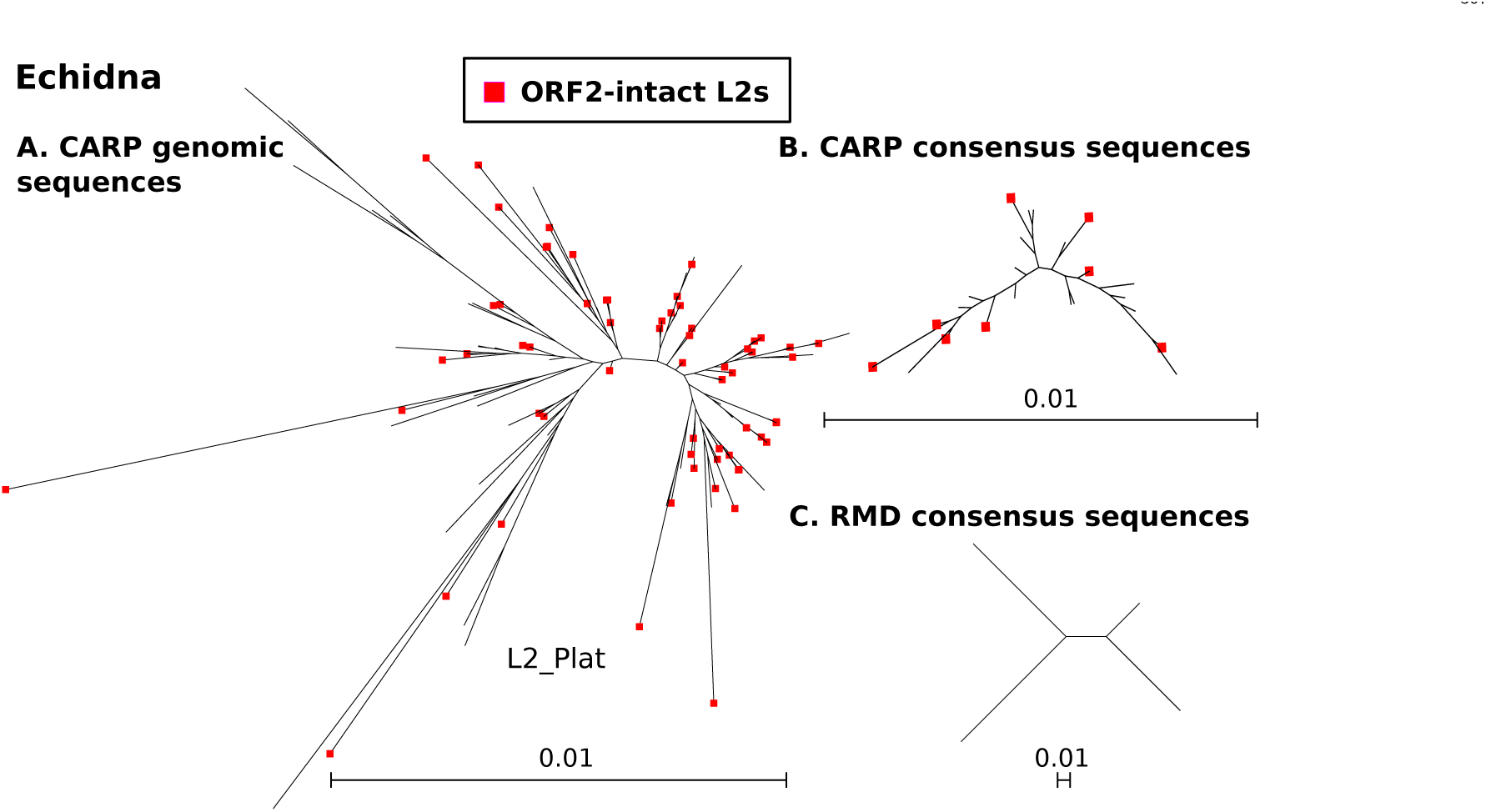
Phylogenetic analysis of L2 elements in the echidna genome. Figure shows the dendrograms of full-length L2 elements in the echidna genome. Panel A) long L2 sequences from the echidna genome. Panel B) Long L2 CARP consensus sequences from echidna. Panel C) Long L2 RMD consensus sequences from echidna. Sequences were aligned with MUSCLE, trees inferred with FastTree and visualized with Archaeopteryx. ORF2-instact L2s are shown with a red dot at the tip of the branch.

It is worth noting that the Repbase annotation for L2s puts the full-length platypus L2 consensus sequences at 5kb long. However, based on both the CARP and RMD identification outputs, L2 elements in echidna and platypus were significantly shorter, at 3kb, with the longest one we could find (in the platypus genome) 3,110bp in length.

## Discussion

Design considerations for bioinformatics pipelines or packages to identify and annotate repetitive sequences in DNA reflect the (sometimes unstated) goals of their developers. We have chosen to prioritise the identification of recent, slightly divergent repeats, to do so with a minimal number of tools and dependencies and to allow users the flexibility of choosing their own annotation tools (ie RepeatMasker or CENSOR). Ancestral, or previously characterised repeats can easily be detected using existing tools, but identifying novel repetitive elements, such as clade specific SINEs requires *ab initio* identification. It is also our experience that researchers sequencing a new genome usually want to identify repeats and segmental duplications early, and independently of gene model prediction. CARP is based on PALS and PILER, but improved and re-implemented in Go as krishna and igor. Our pipeline boils down to five simple steps: 1) find repeats using a pair-wise all vs all local alignment in your genome of choice, 2) use single linkage clustering with a length constraint to create repeat families from the alignments, 3) generate consensus sequences from repeat families and annotate them using RepBase and reverse transcriptase sequences and TE sequences from NCBI, 4) filter out protein coding genes by alignment to Swiss-Prot and 5) combine the *ab initio* library with RepBase to annotate both TEs and candidate segmental duplications.

At present RMD is the most widely used *ab initio* TE identification package, but it has limitations, particularly for users interested in the evolution of TEs. It only provides broad consensus sequences and does not allow one to determine what sequences contributed to a consensus. REPET can provide the sequences/genome intervals used to generate the consensus sequences from PILER families, but not for GROUPER or RECON families. Neither RMD nor REPET removes families obtained from gene families as does CARP. REPET will use gene models to filter out gene repeats, but if no gene model intervals are available this is not an option.

Neither RMD nor REPET are designed to detect segmental duplications. REPET in particular is designed to remove low copy number families from the analysis in order to avoid having segmental duplications in the final consensus set. At present, segmental duplications are detected using all vs all pair-wise alignments of TE repeat masked genomes, because TE repeats generate a huge number of alignments that mask the *bona fide* segmental duplications. This masking of TEs also reduces the sensitivity of existing segmental duplication approaches as TEs are a significant component of segmental duplication sequences. CARP generates consensus sequences from low copy repeats (segmental duplications) without masking, which improves the sensitivity of segmental duplication detection. When we compared our segmental duplication annotation to what has been reported for these seven species, we found that our method detected more candidate segmental duplications in the anolis (4.9%) [40] (Table 3) and the opossum (1.7%) (Table 3) [41].

Finally, the echidna genome is made up of almost 24% LINE L2 sequences, which is an extraordinarily high percentage. Such a high percentage of a single repeat type usually means that there are many actively retrotransposing elements in the genome. As part of CARP’s standard output, we were able to identify 110 potentially active, L2 elements in the echidna genome with minimal additional analysis (Figure 4).

### Conclusion

Here we introduce a simple and flexible *ab initio* repeat identification and annotation method (CARP) that annotates TEs and candidate segmental duplications. We applied CARP to seven animal genomes and demonstrated that it performs as well or better than RepeatModeller, the most commonly used *ab initio* TE annotation package.

Limitation: Our approach is limited by memory requirements and runtime. However, as hardware improves and becomes less expensive, these limitations will become less of an issue.

## Supporting information

**S1 Fig. Coverage plot of high copy number unclassified repeats in the anole genome.** Shows the top 12 highest copy number (>2,000 copies) unclassified consensus sequences coverage plot in the anole genome.

**S2 Fig. Coverage plot of high copy number unclassified repeats in the opossum genome.** Shows the top 5 highest copy number (>2,000 copies) unclassified consensus sequence coverage plots in the opossum genome.

**S3 Fig. Coverage plot of high copy number unclassified repeats in the human genome.** Shows the top 5 highest copy number (>2,000 copies) unclassified consensus sequences coverage plot in the human genome.

**S1 Appendix. CARP documentation.** Gives a detailed account of how to use our *ab initio* method to identify and annotate TEs from a genome assembly, including the benchmarks used for the seven species in this report.

**S1 Table. Genome dataset.** Shows the systematic name, common name, genome version, source and submitter for all the genomes tested for our *ab initio* method. Genomes that were acquired through private collaboration (not publicly avaiable) are marked as ‘Private’ in the source column.

**S2 Table. Assembly statistics.** Shows the systematic name, total sequence length, scaffold N50, contig N50 and assembly level.

**S3 Table. Assembly method and coverage.** Shows the systematic name, assembly method, sequencing techonology and estimated genome coverage for the seven genomes in this study.

**S4 Table. Bechmarks for each methods.** Here we show the compute time used for the seven tested species with CARP and RMD.

**S5-11 Table. Repeat content in seven target species.** Here we show the proportion and copy number of each repeat class in chicken, anole, bearded dragon, opossum, platypus, echidna and human.

## Acknowledgments

We would like to thank our colleagues (Frank Grützer from the University of Adelaide and Guojie Zhang, University of Copenhagen) for making the private echidna genome assembly available to us. We would also like thank James Galbraith for taking the time to read the manuscript in full and offering helpful comments. Finally, this paper would not be possible without the invaluable insights and writing help from Atma Ivancevic, brainstorming from Reuben Buckley, and extraodinary IT support from Matt Westlake.

## Availability and requirements

The Go source code are available on github: github.com/biogo/examples/krishna

## References

1. Cordaux R, Batzer MA. The impact of retrotransposons on human genome evolution. Nature reviews Genetics. 2009;10(10):691.

2. Schnable PS, Ware D, Fulton RS, Stein JC, Wei F, Pasternak S, et al. The B73 maize genome: complexity, diversity, and dynamics. science. 2009;326(5956):1112–1115.

3. Lander ES, Linton LM, Birren B, Nusbaum C, Zody MC, Baldwin J, et al. Initial sequencing and analysis of the human genome. 2001;.

4. Eichler EE. Recent duplication, domain accretion and the dynamic mutation of the human genome. TRENDS in Genetics. 2001;17(11):661–669.

5. Muñoz-López M, García-Pérez JL. DNA transposons: nature and applications in genomics. Current genomics. 2010;11(2):115–128.

6. Skipper KA, Andersen PR, Sharma N, Mikkelsen JG. DNA transposon-based gene vehicles-scenes from an evolutionary drive. Journal of biomedical science. 2013;20(1):92.

7. Khodosevich K, Lebedev Y, Sverdlov E. Endogenous retroviruses and human evolution. Comparative and functional genomics. 2002;3(6):494–498.

8. Maksakova IA, Romanish MT, Gagnier L, Dunn CA, Van de Lagemaat LN, Mager DL. Retroviral elements and their hosts: insertional mutagenesis in the mouse germ line. PLoS genetics. 2006;2(1):e2.

9. Lee SI, Kim NS. Transposable elements and genome size variations in plants. Genomics & informatics. 2014;12(3):87–97.

10. Holton NJ, Goodwin TJ, Butler MI, Poulter RT. An active retrotransposon in Candida albicans. Nucleic acids research. 2001;29(19):4014–4024.

11. Matthews GD, Goodwin T, Butler MI, Berryman TA, Poulter R. pCal, a highly unusual Ty1/copia retrotransposon from the pathogenic yeast Candida albicans. Journal of bacteriology. 1997;179(22):7118–7128.

12. Nefedova L, Kim A. Molecular phylogeny and systematics of Drosophila retrotransposons and retroviruses. Molecular biology. 2009;43(5):747.

13. Luan DD, Korman MH, Jakubczak JL, Eickbush TH. Reverse transcription of R2Bm RNA is primed by a nick at the chromosomal target site: a mechanism for non-LTR retrotransposition. Cell. 1993;72(4):595–605.

14. Ivancevic AM, Kortschak RD, Bertozzi T, Adelson DL. LINEs between Species: Evolutionary Dynamics of LINE-1 Retrotransposons across the Eukaryotic Tree of Life. Genome Biol Evol. 2016;8(11):3301–3322. doi:10.1093/gbe/evw243.

15. Smit AFA, Hubley R, Green P. RepeatMasker Open-4.0.; 2013-2015.

16. Jurka J, Kapitonov VV, Pavlicek A, Klonowski P, Kohany O, Walichiewicz J. Repbase Update, a database of eukaryotic repetitive elements. Cytogenetic and genome research. 2005;110(1-4):462–467.

17. McCarthy EM, McDonald JF. LTR_STRUC: a novel search and identification program for LTR retrotransposons. Bioinformatics. 2003;19(3):362–367.

18. Flutre T, Duprat E, Feuillet C, Quesneville H. Considering transposable element diversification in de novo annotation approaches. PLoS One. 2011;6(1):e16526. doi:10.1371/journal.pone.0016526.

19. Girgis HZ. Red: an intelligent, rapid, accurate tool for detecting repeats de-novo on the genomic scale. BMC bioinformatics. 2015;16(1):227.

20. Edgar RC, Myers EW. PILER: identification and classification of genomic repeats. Bioinformatics. 2005;21(suppl_1):i152–i158.

21. Price AL, Jones NC, Pevzner PA. De novo identification of repeat families in large genomes. Bioinformatics. 2005;21(suppl_1):i351–i358.

22. Bao Z, Eddy SR. Automated de novo identification of repeat sequence families in sequenced genomes. Genome research. 2002;12(8):1269–1276.

23. Benson G. Tandem repeats finder: a program to analyze DNA sequences. Nucleic acids research. 1999;27(2):573.

24. Quesneville H, Nouaud D, Anxolabehere D. Detection of new transposable element families in Drosophila melanogaster and Anopheles gambiae genomes. J Mol Evol. 2003;57 Suppl 1:S50–9. doi:10.1007/s00239-003-0007-2.

25. Quesneville H, Bergman CM, Andrieu O, Autard D, Nouaud D, Ashburner M, et al. Combined evidence annotation of transposable elements in genome sequences. PLoS Comput Biol. 2005;1(2):166–275. doi:10.1371/journal.pcbi.0010022.

26. Kortschak RD, Adelson DL. biogo: a simple high-performance bioinformatics toolkit for the Go language. bioRxiv. 2014; p. 005033.

27. Edgar RC. MUSCLE: multiple sequence alignment with high accuracy and high throughput. Nucleic acids research. 2004;32(5):1792–1797.

28. Gish W. Wu-blast; 1996.

29. Kohany O, Gentles AJ, Hankus L, Jurka J. Annotation, submission and screening of repetitive elements in Repbase: RepbaseSubmitter and Censor. BMC bioinformatics. 2006;7(1):474.

30. Pruitt KD, Tatusova T, Maglott DR. NCBI reference sequences (RefSeq): a curated non-redundant sequence database of genomes, transcripts and proteins. Nucleic acids research. 2006;35(suppl_1):D61–D65.

31. Boeckmann B, Bairoch A, Apweiler R, Blatter MC, Estreicher A, Gasteiger E, et al. The SWISS-PROT protein knowledgebase and its supplement TrEMBL in 2003. Nucleic acids research. 2003;31(1):365–370.

32. Quinlan AR, Hall IM. BEDTools: a flexible suite of utilities for comparing genomic features. Bioinformatics. 2010;26(6):841–842.

33. Castresana J. Selection of conserved blocks from multiple alignments for their use in phylogenetic analysis. Molecular biology and evolution. 2000;17(4):540–552.

34. Price MN, Dehal PS, Arkin AP. FastTree: computing large minimum evolution trees with profiles instead of a distance matrix. Molecular biology and evolution. 2009;26(7):1641–1650.

35. Edgar RC. Search and clustering orders of magnitude faster than BLAST. Bioinformatics. 2010;26(19):2460–2461.

36. Finn RD, Clements J, Eddy SR. HMMER web server: interactive sequence similarity searching. Nucleic acids research. 2011;39(suppl_2):W29–W37.

37. Bateman A, Coin L, Durbin R, Finn RD, Hollich V, Griffiths-Jones S, et al. The Pfam protein families database. Nucleic acids research. 2004;32(suppl_1):D138–D141.

38. Heras S, Thomas M, Garcia-Canadas M, De Felipe P, Garcia-Perez J, Ryan M, et al. L1Tc non-LTR retrotransposons from Trypanosoma cruzi contain a functional viral-like self-cleaving 2A sequence in frame with the active proteins they encode. Cellular and Molecular Life Sciences CMLS. 2006;63(12):1449–1460.

39. Dewannieux M, Esnault C, Heidmann T. LINE-mediated retrotransposition of marked Alu sequences. Nature genetics. 2003;35(1):41.

40. Alfoldi J, Di Palma F, Grabherr M, Williams C, Kong L, Mauceli E, et al. The genome of the green anole lizard and a comparative analysis with birds and mammals. Nature. 2011;477(7366):587.

41. Samollow PB. The opossum genome: insights and opportunities from an alternative mammal. Genome research. 2008;18(8):1199–1215.

